# Ribosomal Remodeling and Translational Suppression in the Post-Arrest Heart

**DOI:** 10.64898/2026.11.05.764511

**Authors:** Yong Gao, William Danysh, Gao Zhou, Kenneth R. Laurita, Lance Wilson, Zhenxiang Gao, Rong Xu, Cody Rutledge

## Abstract

**Background:** Cardiac arrest (CA) is a leading cause of death and disability in the United States, and no pharmacologic therapies improve post-arrest outcomes. Acute myocardial dysfunction is a highly prevalent CA complication that contributes to post-arrest mortality, yet the molecular mechanisms that drive this dysfunction remain incompletely understood. Prior work in our laboratory has suggested that ribosome-associated pathways are altered after CA in the heart and that translational dysregulation may be a central feature of post-arrest myocardial injury.

**Objectives:** To define the impact of CA on myocardial ribosome biology and protein synthesis across experimental models of CA and determine how transcriptome, ribosome-bound transcriptome, translational efficiency, and proteome changes evolve after CA in the heart.

**Methods:** We studied complementary porcine and murine models of CA. In pigs, left ventricular myocardium was collected from naïve controls, cardiac arrest with return of spontaneous circulation (Arrest), and cardiac arrest with pulseless electrical activity (PEA) re-arrest in a myocardial infarction-induced CA model and analyzed tissue by RNA sequencing. In mice, left ventricular tissue was collected at various time points after potassium chloride-induced asystole and resuscitation for ribosome activity and multi-omic profiling including RNA sequencing, ribosome profiling, and proteomics.

**Results:** In the porcine model, myocardial transcriptomes demonstrate enrichment of cytosolic ribosomal protein transcripts and translation-related pathways that increase with ischemic burden. In contrast, a functional assay of ribosome activity in mice demonstrates suppression of protein synthesis after CA with gradual recovery over days. Multi-omic profiling in the mouse model demonstrates that ribosome- and translation-related pathways are among the most consistently altered programs across time points and modalities. While 6 h studies note prevalent inflammatory signals, 24 h studies find disruptions to ion homeostasis and conduction-related pathways that are likely related to translational suppression.

**Conclusions:** CA induces marked remodeling of myocardial ribosome biology, characterized by increasing ribosomal transcripts with ischemic burden in the pig model. However, functional studies in the mouse model demonstrate that ribosome activity is suppressed following CA and remains suppressed for at least 24 h after resuscitation, contributing to reduced nascent protein synthesis and may limit critical functions of the heart. These findings identify translational control as a central feature of post-arrest myocardial injury and suggest that ribosome-dependent recovery pathways may represent a novel therapeutic target in post-arrest cardiac dysfunction.

## Introduction

Cardiac arrest (CA) remains a leading cause of death in the United States, affecting more than 350,000 individuals outside of the hospital every year^1–3^. Despite improvements in bystander cardiopulmonary resuscitation (CPR), emergency medical response, and post-resuscitation critical care, survival after CA remains poor, with out-of-hospital CA survival rates around 10%^1,2^. Many patients who initially achieve return of spontaneous circulation (ROSC) ultimately die from post-arrest complications^2,4^.

Post-CA complications, which are largely attributed to ischemia/reperfusion injury (IRI), can affect nearly every organ in the body^5,6^. While neurologic injury accounts for the majority of post-CA deaths^4^, acute myocardial dysfunction is also highly prevalent and a major contributor to mortality^7^. Up to two-thirds of CA survivors demonstrate systolic and diastolic cardiac dysfunction shortly after ROSC^8,9^. This post-arrest myocardial dysfunction, often termed myocardial stunning, begins immediately after resuscitation and may persist for days^9,10^. The molecular mechanisms driving myocardial stunning remain incompletely understood^11,12^.

Current standards of post-CA care, including temperature control and vasopressor support, address hemodynamic consequences of myocardial dysfunction but do not target its underlying cellular mechanisms^13^. There are no approved pharmacologic therapies that directly attenuate myocardial injury following CA, representing a critical unmet need in the management of post-arrest survivors. Therapies directed at single inflammatory mediators or signaling pathways have not yet demonstrated improved clinical outcomes after CA^12,14,15^. Targeting broader cellular processes that integrate multiple injury pathways may be more effective^16^.

One such mechanism may be disruption of ribosome activity and translational control after IRI. Suppression of protein synthesis is a recognized response to ischemic and hypoxic stress across multiple tissues and is thought to be an adaptive mechanism to conserve energy^17–19^. This suppression is particularly pronounced in tissues with high aerobic demand, such as neurons and cardiomyocytes^20,21^. However, translational suppression that persists after reperfusion may be maladaptive. In the brain, protein synthesis remains suppressed for days after IRI, and failure to restore translation precedes delayed neuronal death in vulnerable regions^18,22,23^. Sustained translational suppression may be particularly consequential in the heart, where continuous protein synthesis is required to maintain the contractile apparatus, ion channel expression, and mitochondrial function^24–26^.

Recent work in our laboratory has found that ribosome activity is reduced in the mouse heart after CA and in cells exposed to hypoxia/reperfusion, and that metformin pretreatment partially preserves ribosome function^27^. However, the extent to which ribosome-associated changes occur across models of CA, how these changes relate to ischemic burden and timing after recovery, and the consequences of reduced ribosome activity for specific proteins and pathways remain unclear.

In this study, we used complementary porcine and murine models of CA to define post-arrest myocardial ribosome biology. The porcine model of myocardial infarction-induced ventricular fibrillation reproduces clinical CA and allows comparison across CA subtypes, while the murine model permits survival time courses and *in vivo* assessment of nascent proteins. We hypothesized that CA induces sustained suppression of myocardial translation that persists after reperfusion across models. We first assessed transcriptome changes in porcine myocardium after CA with ROSC and after CA complicated by PEA re-arrest. We then measured myocardial protein synthesis by puromycin incorporation in mice 6, 24, and 72 h after CA to define the time course of translational recovery. Finally, we performed paired RNA sequencing, ribosome profiling, and proteomics in murine myocardium to determine transcript abundance, translational efficiency (TE), and protein abundance over time after CA. Together, these studies identify ribosome remodeling and translational suppression as central features of the post-arrest myocardial response.

## Materials and Methods

### Pig Model of Cardiac Arrest

An established porcine model of CA was utilized as previously described^28,29^. Briefly, pigs were anesthetized with telazol (2.0 mg/kg), ketamine (1.5 mg/kg) and xylazine 1.5 mg/kg, then maintained on isoflurane (1-3%), mechanically ventilated, and instrumented for continuous monitoring of temperature, heart rate, central arterial pressure, end-tidal CO₂, and oxygen saturation. To create an acute MI and induce ventricular fibrillation (VF), the left anterior descending (LAD) coronary artery was reversibly occluded using a balloon catheter distal to the second diagonal branch. Hemodynamic measurements were recorded at baseline and every 2 min during ischemia and after ROSC, and at all events, including arrhythmia, defibrillation, and drug administration. If spontaneous VF did not occur within 45 min of LAD occlusion, VF was induced by direct current through an electrophysiological catheter placed in the LV. VF was then left untreated for 8 min to simulate out-of-hospital CA, after which animals were resuscitated according to advanced cardiac life support guidelines. Epinephrine (0.01 mg/kg IV) was given every two min during CPR. ROSC was defined as an organized rhythm with MAP >30 mm Hg lasting for ≥1 minute. Re-arrest by VF or pulseless electrical activity (PEA, defined as mean arterial pressure ≤25% of baseline mmHg with an organized rhythm), was treated by resuming ACLS. After 75 min of total LAD occlusion, the LAD was reperfused, and after a total duration of 180 min from LAD occlusion, remote LV myocardium was collected. Animals were classified as Arrest (ROSC without subsequent PEA) or PEA (≥1 episode of PEA re-arrest). Naïve controls underwent the same instrumentation without LAD occlusion or arrest. Animals without ROSC after 20 min of resuscitation and animals with recurrent VF but no evidence of PEA were excluded from analysis.

All pig experiments were performed at MetroHealth Medical Center in compliance with National Institutes of Health guidelines for the care and use of laboratory animals and were approved by the CWRU Institutional Animal Care and Use Committee.

### Pig Transcriptome Data

Left ventricular (LV) biopsies were taken from remote tissue away from the infarct zone 3 h after LAD occlusion, then flash frozen and powdered. Total RNA was isolated from ∼50 mg of tissue using RNeasy kit (Qiagen, Germantown, MD, USA) and sent to Novogene (Sacramento, CA, USA) for commercial RNA sequencing. Sample quality was assessed by Agilent 2100 bioanalyzer (Santa Clara, CA, USA). Samples underwent poly(A) enrichment, RNA fragmentation, reverse transcription of cDNA, and mRNA library preparation according to Novogene specifications. Samples were sequenced on an Illumina NovaSeq platform (San Diego, CA, USA).

Raw sequencing reads were quality controlled for Phred score and GC-content distribution. Reads were aligned with HISAT2 to the pig genome (Sscrofa11.1), and gene-level counts were generated. Low-count genes were filtered before downstream analyses. Differential expression was tested using DESeq2 with batch number and RNA integrity number (RIN) included as covariates. Pre-ranked gene set enrichment analysis (GSEA) was performed using fgsea with Reactome pathway collections as described below. Cytosolic ribosomal protein genes were additionally analyzed as a focused gene set across comparisons. Sixteen animals were included in the final analysis: Naïve (n = 6), Arrest (n = 4), and PEA (n = 6). Two samples with low RNA integrity (RIN 4.1 and 4.8) were removed from the arrest group. Ribosomal protein (RP) score was defined as the mean z-score of Rpl/Rps families, excluding mitochondrial Mrpl/Mrps genes and pseudogenes, after adjusting for RIN (Supplemental Figure 2B) and was compared by one-way ANOVA. The relationship between RP reads and RIN was assessed by Spearman correlation (Supplemental Figure 2C).

### Mouse Model of Cardiac Arrest

Eight- to twelve-week-old male and female C57BL/6J mice (Jackson Labs, Bar Harbor, ME, USA) were studied. Mice were randomly divided into Naïve groups (echocardiography only) or CA procedure, as previously described^30^. Briefly, mice were anesthetized using 5% vaporized isoflurane in an induction chamber, followed by rapid endotracheal intubation using a 20-gauge catheter. Mice were placed onto a preclinical echocardiography platform (Prospect T1, Scintica, Webster, TX, USA) with ECG tracing and heating pad with rectal temperature probe. Mice were mechanically ventilated using a Minivent (Harvard Apparatus, Holliston, MA, USA) system at 150 breaths/min x 125 µL (females) or 140 µL (males). Isoflurane was titrated between 1-2% to maintain heart rate at 400-500 bpm pre-procedurally, body temperature was maintained at 36-37°C, and baseline echocardiography obtained.

After depilation and sterile preparation, a sterile 32-gauge needle was guided apically into the LV cavity under ultrasound guidance. 40 µL of 0.5 M KCl was delivered into the LV, causing immediate asystole. Ventilation and isoflurane were stopped. Mice remained in asystole for 8 min. Immediately before the 8-min mark, 500 µL of 15 µg/mL epinephrine in normal saline was injected into the LV cavity and cardiac resuscitation (300 bpm by finger compressions) began. If a ventricular rhythm was observed on ECG, compressions were stopped and Doppler echocardiography was used to confirm ROSC. If ROSC was not obtained within 3 min, the procedure was abandoned. After achieving ROSC, mice were maintained on ventilation until spontaneous respiration exceeded 60 breaths/min, prompting extubation and transfer to a heated recovery cage.

Mice were monitored in recovery cages for 6, 24, or 72 h. Terminal cardiac contractility and a 12-point neurologic exam were performed as previously described^31,32^. At terminal time points, animals were euthanized following imaging and hearts rapidly excised into cold saline, where LVs were dissected and then flash frozen. The apex of the LV was discarded to avoid needle penetration injury. All mouse experiments were performed at the Louis Stokes Cleveland VA Medical Center in compliance with National Institutes of Health Guidance for Care and Use of Experimental Animals and were approved by the VA Institutional Animal Care and Use Committee.

### Puromycin Incorporation and Western Blot Analysis

One hour before euthanasia, a cohort of mice (n=6 per group) from Naïve, 6, 24, and 72 h CA cohorts received intraperitoneal injection of puromycin (0.04 µmol/g body weight; approximately 22 mg/kg) for puromycin incorporation study of ribosome activity^33^. Tissues were flash frozen for western blot analysis. Frozen tissue was homogenized in lysis buffer containing protease inhibitor and normalized for protein concentration. Samples were heated and separated by SDS-PAGE on 4-12% gradient gels, then transferred to nitrocellulose membrane and immediately stained with ponceau solution as a loading control. The ponceau stain was washed away and membrane blocked with 5% milk in TBS-tween for 1 h, then incubated overnight at 4°C with puromycin antibody (Developmental Studies Hybridoma Bank, University of Iowa, USA, #PMY-2A4, 1:1000). Following incubation, membranes were washed and then probed for 1 h with HRP-conjugated anti-mouse IgG2 secondary and images acquired using chemiluminescent detection. Images were analyzed using Fiji^34^. Total lane puromycin signal was normalized to total lane Ponceau signal and expressed relative to the Naïve mean.

### Mouse Multi-omics Data

LV tissue was collected from Naïve, 6, and 24 h CA cohorts (n=6/group, 3 male and 3 female) for multi-omic profiling. Tissues were frozen and powdered and divided for RNA-sequencing/ribosome sequencing and proteomics.

RNA sequencing and ribosome profiling were performed by the CWRU Advanced RNA Profiling Core as previously described^35^. Briefly, powdered LV tissue was lysed in buffer containing cycloheximide, and each lysate was divided for RNA sequencing and ribosome profiling. For ribosome profiling, lysates were nuclease digested and monosomes were isolated by sucrose gradient fractionation. Ribosome-protected fragments (RPFs) were size-selected by urea-PAGE and depleted of ribosomal RNA. RNA-seq libraries were prepared from total RNA. Libraries were sequenced on an Illumina NextSeq 550. Gene-level counts were generated over exons for RNA-seq and over coding sequences for RPFs.

For RNA-seq and ribosome profiling, genes with at least 10 reads in at least as many samples as the smallest group were retained and differential expression was tested with DESeq2 1.52.0 using the design ∼ sex + batch + condition with local dispersion estimation. Each arrest timepoint was compared with Naive controls by Wald test, with Benjamini-Hochberg adjustment and an adjusted p value < 0.05 considered significant. Translational efficiency, defined for each gene as the ratio of ribosome-protected fragment (RPF) to RNA abundance, was assessed in two ways. Gene-level changes were tested by the deltaTE approach, in which RNA and ribosome-protected fragment counts were modeled jointly (∼ sex + batch + condition + assay + condition:assay) and the condition-by-assay interaction term was tested. Global translational efficiency was summarized by regressing ribosome-protected fragment log2 fold changes on RNA log2 fold changes, where a slope below unity indicates that ribosome occupancy does not scale with transcript abundance. Per-gene translational efficiency was additionally calculated as log2 of the ratio of mean ribosome-protected fragment counts per million to mean RNA counts per million within each condition and displayed as arrest versus Naive. One 24 h ribosome profiling sample was excluded for insufficient depth (0.06 million assigned reads, 1,440 genes detected), leaving n = 5 at 24 h for all sequencing analyses.

Proteomic studies were completed by MS Bioworks (Ann Arbor, MI, USA). Tissue was lysed in SDS-containing RIPA buffer, TCA precipitated, and resuspended in 8 M urea. Protein was reduced, alkylated, and digested overnight with trypsin, then desalted and peptides analyzed by data-independent acquisition on an Orbitrap Astral mass spectrometer (Thermo Fisher) coupled to a Vanquish Neo UPLC with a 30 min gradient. A pooled reference sample was injected at the start and end of the batch to assess reproducibility. Data were processed in DIA-NN (v1.9.2) against the SwissProt mouse database at 1% peptide and protein FDR, and normalized intensities were log2 transformed. Two 6 h proteomics samples were excluded by the proteomics core for low protein yield, leaving n = 4 for the 6 h group. Proteins quantified in every sample were tested with limma using the design ∼ sex + condition, with empirical Bayes moderation and Benjamini-Hochberg adjustment. For visualization, counts were variance-stabilized and the sex and batch terms were removed by linear regression with condition retained in the model; the same adjustment was applied to the proteomics matrix. Principal component analysis and hierarchical clustering (1 − Pearson correlation, average linkage) used the 500 most variable features. Heatmaps display row-standardized adjusted values truncated at ±2.

Pre-ranked gene set enrichment analysis was performed for each modality and timepoint with fgsea 1.38.0, ranking genes by the sign of the log2 fold change multiplied by −log10(p value). Gene sets were obtained from the Molecular Signatures Database through msigdbr 26.1.0 and restricted to Reactome collections. Gene sets containing 15 to 500 genes were tested using the multilevel algorithm with Benjamini-Hochberg adjustment within each collection, and redundant sets were collapsed with fgsea’s collapsePathways. Cytosolic ribosome and translation gene sets, together with a set comprising the detected cytosolic ribosomal protein genes, were tested across all modalities and timepoints as a focused panel. Figures were generated with ggplot2 4.0.3 and pheatmap 1.0.13.

### Statistical Analysis

Data were expressed as mean ± SEM. One-way ANOVA with Dunnett’s multiple comparisons test was used for multiple-group comparisons. Apart from -omics analyses (detailed above), p < 0.05 was considered significant for all comparisons. All statistical analysis was completed using GraphPad Prism 8.0.5 (San Diego, CA, USA) software or R 4.6.1.

### Statement of data availability

mRNA sequencing and ribosequencing data has been uploaded to the NIH GEO database (accession #GSE353360 and #GSE353540). Raw data are available from the corresponding author upon reasonable request.

## Results

### Ribosome-associated transcripts increase with ischemic burden in porcine myocardium after arrest

To determine whether ribosome-related programs are altered across increasing degrees of myocardial ischemic stress, we first analyzed LV transcriptomes from a porcine model including Naïve controls, animals with CA with ROSC (Arrest), and animals with CA complicated by PEA (Table 1). Tissue was collected from remote myocardium after reperfusion to assess global transcriptional responses associated with arrest severity (Figure 1A). Principal component analysis demonstrated clear group separation, indicating distinct myocardial transcriptional states between arrest types (Figure 1B). Relative to Naïve myocardium, Arrest animals showed broad transcriptomic remodeling, with 545 genes increased and 352 genes decreased. PEA animals demonstrated 2,542 genes increased and 2,004 genes decreased relative to naïve controls (Figure 1C, D). A prominent feature of both comparisons was the coordinated enrichment of cytosolic ribosomal protein and translation associated transcripts. Cytosolic ribosomal protein genes were particularly enriched in PEA animals (Figure 1D, highlighted in orange), which features a higher ischemic burden than Arrest animals (Table 1).

**Figure 1.**
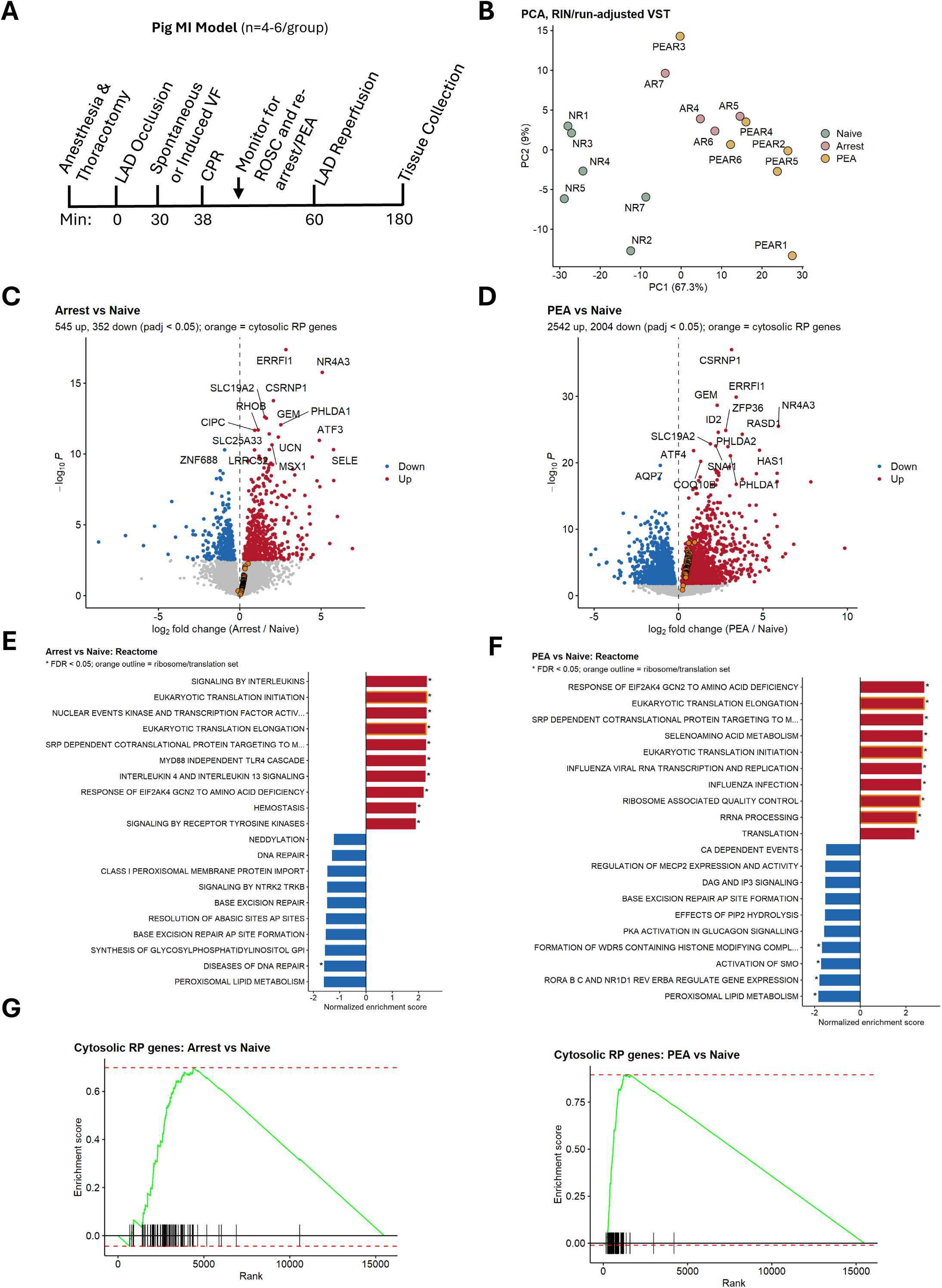
Ribosomal transcripts increase with ischemic burden in porcine remote myocardium after cardiac arrest. A) Study overview depicting LAD occlusion, ventricular arrhythmia, resuscitation, and reperfusion, with timing of tissue collection. B) Principal component analysis of remote myocardial transcriptomes in Naïve, Arrest (ROSC without rearrest), and PEA (≥1 PEA rearrest) animals. C–D) Volcano plots comparing Arrest (C) and PEA (D) animals to Naïve controls; cytosolic ribosomal protein (RP) genes are highlighted in orange. E–F) Top 10 up- and downregulated Reactome pathways by gene set enrichment analysis for Arrest (E) and PEA (F) versus Naïve; ribosome and translation pathways are outlined in orange. G) Enrichment plots for the 78 detected cytosolic RP genes in Arrest versus Naïve (left) and PEA versus Naïve (right). n = 6 Naïve, 4 Arrest, 6 PEA. Differential expression by DESeq2; adjusted p < 0.05 by Benjamini-Hochberg. *=FDR < 0.05 for pathway enrichment.

**Table 1.** Pig Physiologic Data. Values in parentheses are total animals, including those excluded for low RNA integrity. Naïve pigs did not undergo cardiac arrest. Arrest pigs achieved ROSC without re-arrest. PEA pigs had ≥1 PEA re-arrest after initial ROSC requiring CPR and epinephrine. CPR duration is total CPR time, including CPR for re-arrest. Data expressed as mean ± SEM. *p < 0.05 vs. Naïve (one-way ANOVA with Tukey’s post hoc test); ^†^p < 0.05 vs. arrest (Welch’s t-test). CPR, cardiopulmonary resuscitation; Epi, epinephrine; MAP, mean arterial pressure; PEA, pulseless electrical activity; ROSC, return of spontaneous circulation; VF, ventricular fibrillation.

|  | **Number of Pigs Studied (Total)** | **% Female** | **Weight (kg)** | **VF duration (min)** | **CPR duration (min)** | **Shocks delivered** | **Epi doses** | **Time to ROSC (min)** | **Terminal MAP (mmHg)** |
|---|---|---|---|---|---|---|---|---|---|
| **Naïve** | 6 (6) | 100 | 30.4±1.6 | n/a | n/a | n/a | n/a | n/a | 52.1±3.2 |
| **Arrest** | 4 (6) | 100 | 32.0±2.0 | 4.6±2.6 | 6.5±1.6 | 0.5±0.3 | 2.8±0.5 | 8.3±1.2 | 50.8±4.5 |
| **PEA** | 6 (6) | 100 | 33.4±2.2 | 13.5±2.4<sup>†</sup> | 18.4±5.1 | 3.5±1.0<sup>†</sup> | 5.3±1.3 | 18.7±5.2 | 24.7±8.5* |

GSEA Reactome pathway analysis reinforced this observation, with ribosome- and translation-related pathways among the most strongly enriched programs in both Arrest and PEA myocardium (Figure 1E–F), though inflammatory signals were also highly prevalent in the Arrest cohort. GSEA specific to cytosolic ribosomes confirmed significant enrichment of the detected cytosolic ribosomal protein gene set in both Arrest versus Naïve and PEA versus Naïve comparisons, with the strongest effect seen in PEA animals (Figure 1G).

To directly compare transcriptomic responses across levels of post-arrest severity, we next contrasted PEA animals with Arrest animals (Supplemental Figure 1). Sample correlation and clustering analyses demonstrate group separation between arrest type (Supplemental Figure 1 A, B). Differential expression analysis reveals few transcript differences between PEA and Arrest, however, cytosolic ribosomal protein genes remained enriched among the differentially expressed transcripts (Supplemental Figure 1C). GSEA pathway analysis similarly identified translation-related programs, translational elongation, and ribosome-associated pathways among the most strongly enriched processes in PEA relative to Arrest (Supplemental Figure 1D, E). RP scores, calculated from the 78 detected cytosolic RP transcripts, were higher in PEA animals than in both Naïve (p < 0.0001) and Arrest animals (p < 0.01), with Arrest animals intermediate between Naïve and PEA (p = 0.13 vs Naïve; Supplemental Figure 2A, B). Because RNA degradation can confound analysis of ischemic tissues, we examined whether the ribosome-associated transcriptional signal correlated with RIN. The fraction of reads assigned to cytosolic ribosomal protein genes did not significantly correlate with RIN (Supplemental Figure 2C), suggesting that the ribosome signal was not merely an artifact of RNA quality. Together, these findings indicate that cytosolic ribosomal transcripts increase after CA and increase further in the more severe PEA group.

### Global myocardial protein synthesis is suppressed after cardiac arrest in mice

Given the increased ribosomal transcripts in the porcine model, we next measured myocardial protein synthesis during recovery from CA. Because the porcine model is limited by cost and survival time, we used an established mouse model of KCl-induced asystole and resuscitation^30^, with tissue collected from Naïve animals and at 6, 24, and 72 h after CA (Figure 2A; Table 2).

**Figure 2.**
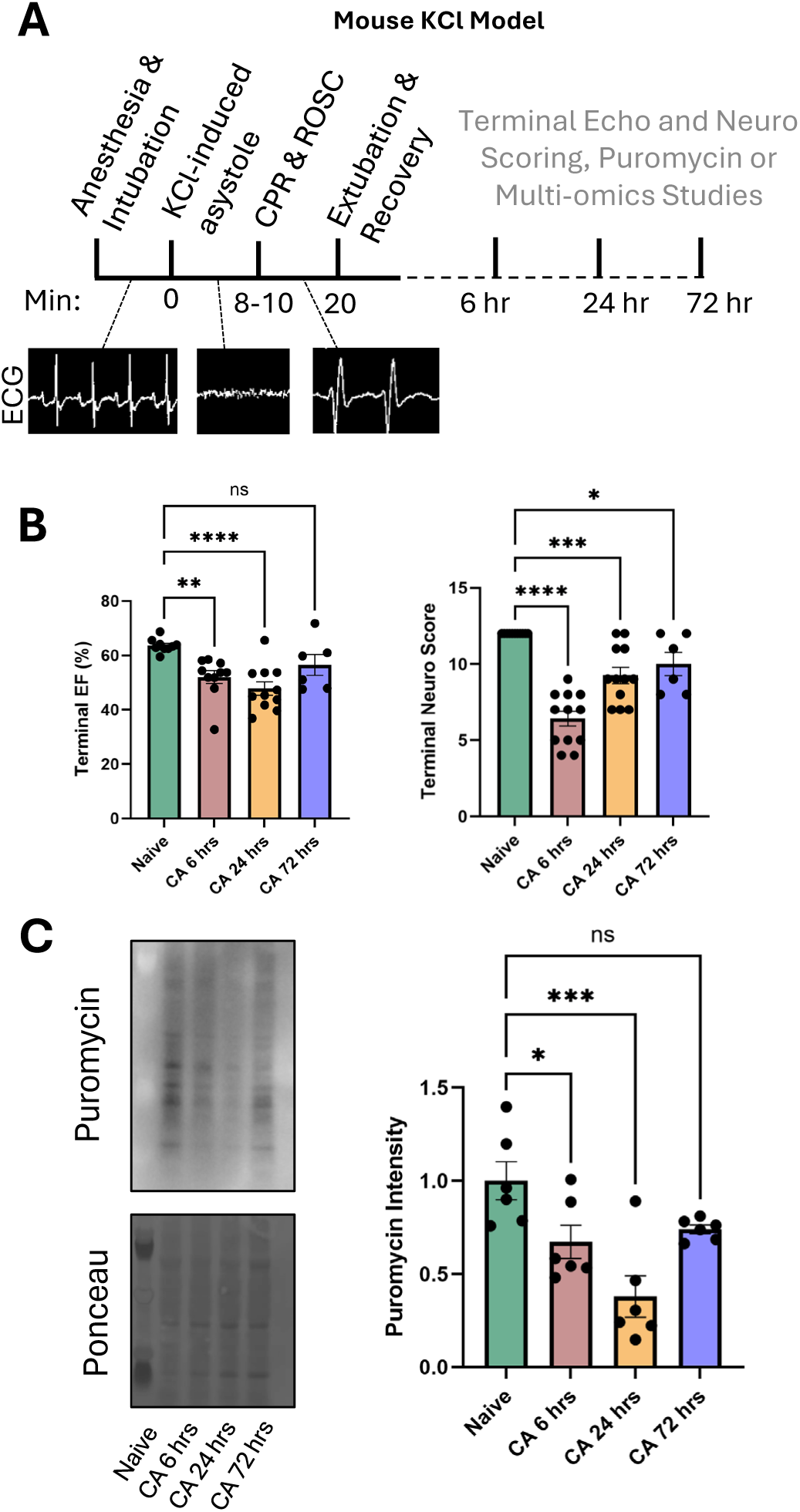
Myocardial protein synthesis is suppressed after cardiac arrest in mice. A) Study overview of the KCl model, depicting anesthesia and intubation, KCl-induced asystole, CPR and ROSC, and extubation and recovery, with representative ECG tracings. Terminal echocardiography, neurologic scoring, and tissue collection for puromycin incorporation or multi-omic studies were performed 6, 24, or 72 h after CA. B) Terminal left ventricular ejection fraction (EF; left) and neurologic score (right) in Naïve and CA animals. C) Representative puromycin immunoblot with Ponceau total protein stain (left) and quantification of puromycin incorporation (right). Puromycin signal was normalized to Ponceau and expressed relative to the Naïve mean. Full blots are shown in Supplemental Figure 3. n = 6-12 mice /group. Data expressed as mean ± SEM. One-way ANOVA with Dunnett’s multiple comparisons test versus Naïve. *p<0.05, **p<0.01, ***p<0.001, ****p<0.0001; ns, not significant.

**Table 2.**
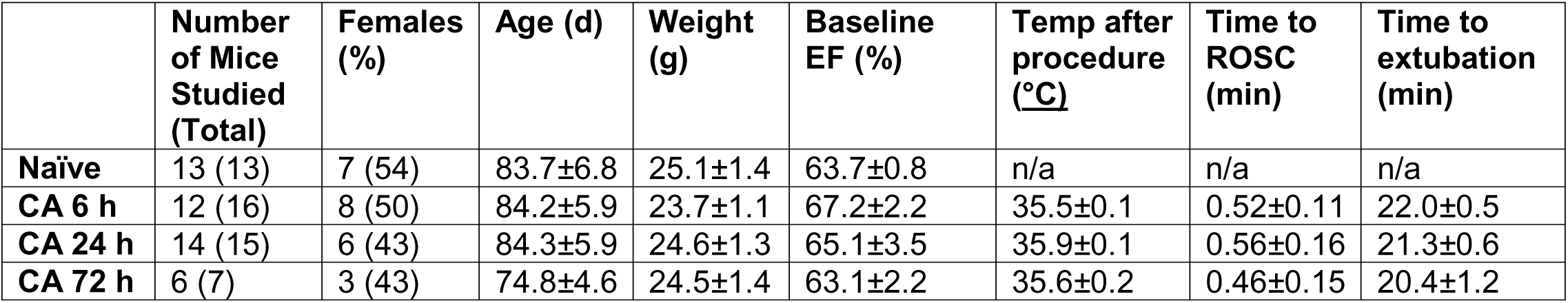
Mouse Physiologic Data. Values in parentheses are total animals, including those excluded for failure to achieve ROSC or death prior to end-point. Females (%) is relative to total animals. Baseline EF was measured before CA. Temperature was measured at the time of extubation. Time to ROSC and extubation were measured from KCl injection. Naïve mice underwent echocardiography only. Data expressed as mean ± SEM. CA, cardiac arrest; EF, ejection fraction; ROSC, return of spontaneous circulation.

CA reduced LV ejection fraction (EF) relative to Naïve (63.7±0.8%) at 6 h (52.0±2.4%, p<0.01), and at 24 h (47.8±2.5%, p<0.0001). EF recovered to 56.5±3.9% at 72 h, which did not differ significantly from Naïve. Neurologic scores (ranked 0-12, where 12 is full neurologic response) were similarly reduced at 6 h (6.4±0.5 vs. 12.0±0.0 in Naïve, p<0.0001) and improved over time, but remained below Naïve at 24 h (9.3±0.5, p<0.001) and 72 h (10.0±0.8, p<0.05; Figure 2B).

Despite the ribosome-associated transcriptional upregulation observed in pigs in both arrest and PEA groups, global protein synthesis in murine myocardium was suppressed after CA, as assessed by reduced puromycin incorporation in left ventricular tissue (Figure 2C; Supplemental Figure 3). Puromycin incorporation was significantly reduced 6 h after CA (67.2±8.9%) compared to Naïve (100±10%, p<0.05), and further reduced at 24 h (37.9±11.1%, p<0.001 vs Naïve) and partially recovered at 72 h (74.0±2.3%, ns vs Naïve), suggesting that translational suppression is an early feature of post-arrest myocardial injury. Although RP transcripts increase in porcine myocardium after CA, global myocardial protein synthesis in mice is suppressed, reaching a nadir at 24 h and no longer differing significantly from Naïve by 72 h.

### Multi-omic profiling reveals distinct changes in transcript abundance, ribosome occupancy, and translational efficiency after arrest

To define the molecular consequences of translational suppression, we performed paired RNA-seq, ribosome profiling (Ribo-seq), and proteomics in murine LV myocardium at 6 h and 24 h after CA (Figure 3A), spanning the early decline and nadir of protein synthesis (Figure 2C).

**Figure 3.**
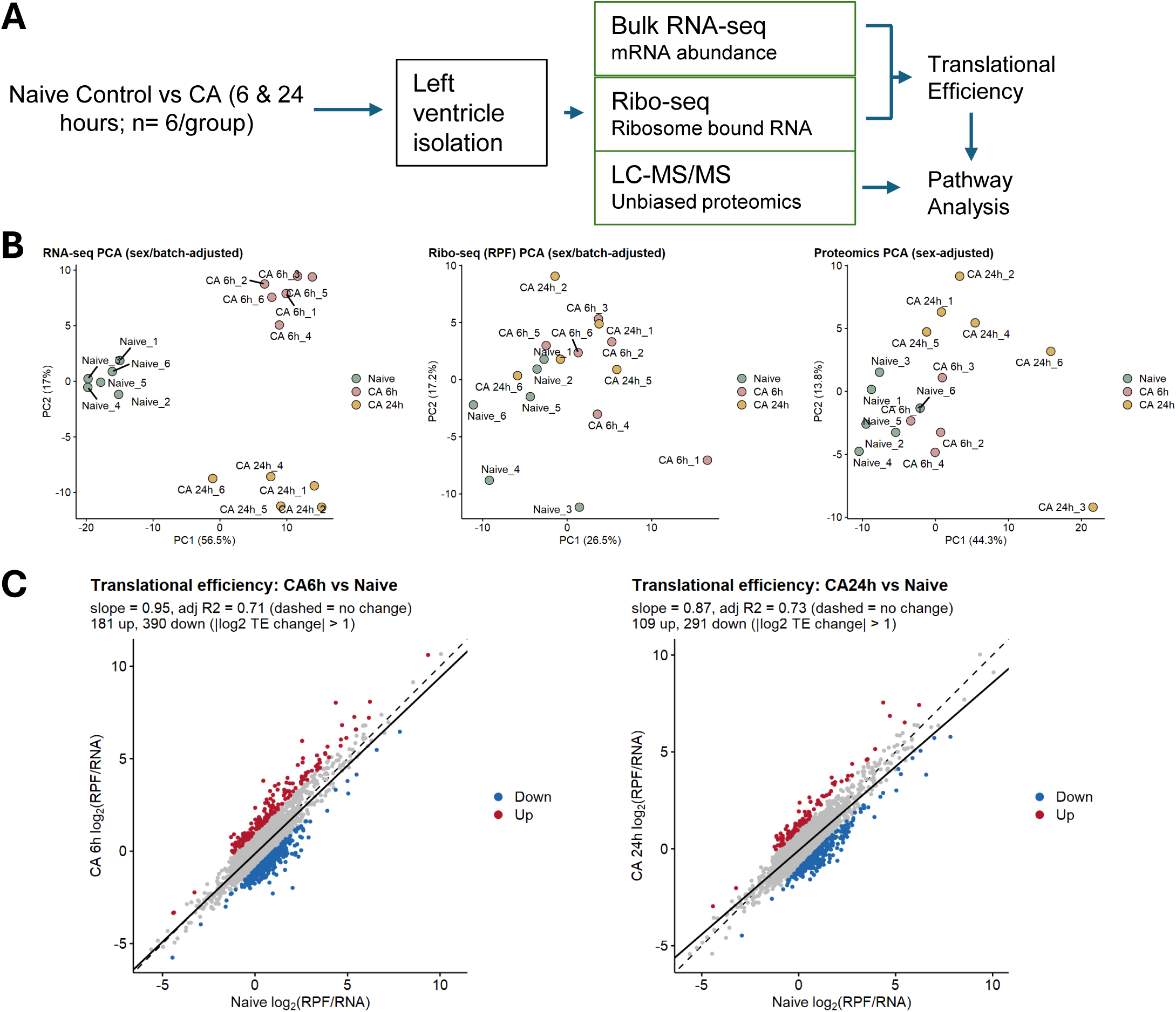
Multi-omic profiling of murine myocardium after cardiac arrest. A) Left ventricular tissue was collected from Naïve mice and 6 h and 24 h after KCl-induced cardiac arrest (CA) for bulk RNA-seq, ribosome profiling (Ribo-seq), and LC-MS/MS proteomics. RNA-seq and Ribo-seq from the same animals were combined to estimate translational efficiency (TE). B) Principal component analysis of RNA-seq (left) and Ribo-seq (center), adjusted for sex and sequencing batch, and of proteomics (right), adjusted for sex. C) Per-gene TE in Naïve versus CA 6h (left) and CA 24h (right) myocardium. TE was calculated as the log2 ratio of mean ribosome-protected fragment (RPF) to mean RNA counts per million in each group. The dashed line indicates unity and the solid line the least-squares fit; slope and adjusted R² are shown. Genes with a greater than 2-fold increase (red) or decrease (blue) in TE relative to Naïve are highlighted. n = 6 Naïve, 6 CA 6h, and 5 CA 24h for sequencing; n = 6 Naïve, 4 CA 6h, and 6 CA 24h for proteomics.

While RNA-seq measures transcript abundance, Ribo-seq quantifies ribosome-protected mRNA fragments (RPFs), providing a transcriptome-wide measure of ribosome engagement on each transcript^36^. Principal component analysis demonstrated separation of Naïve and post-arrest groups within each modality, though most profoundly on an RNA-seq level, indicating robust effects of arrest on transcript abundance, ribosome occupancy, and protein expression (Figure 3B). Sample correlation and heatmap analyses similarly showed reproducible clustering by group across RNA-seq, ribosome profiling, and proteomic datasets (Supplemental Figures 4-6).

We next assessed translational efficiency (TE), defined as RPF abundance relative to RNA abundance for each gene. Per-gene TE after CA remained closely correlated with TE in Naïve myocardium at 6 h (slope 0.95, adjusted R² 0.71) and 24 h (slope 0.87, adjusted R² 0.73; Figure 3C), with modest compression of TE differences between genes at 24 h, despite reduced global protein synthesis (Figure 2C). The relative distribution of ribosomes across transcripts was largely maintained, suggesting that translational suppression after CA is predominantly global rather than transcript-selective.

### Ribosome and translation pathways are among the most consistently altered programs after arrest across modality

We next evaluated differentially expressed genes/proteins and pathway-level changes across modalities. At 6 h after arrest, RNA-seq demonstrated strong induction of inflammatory, stress-response, and cytokine-related pathways together with enrichment of ribosome- and translation-associated pathways, as demonstrated in the pig model (Figure 4A–B). Ribosome profiling at 6 h largely mirrored the inflammatory and stress-response signature, with no significant enrichment of translation pathways and no significantly downregulated pathways (Figure 4C–D). At 24 h after CA, the transcriptional and ribosome occupancy profile shifted. RNA-seq and ribosome profiling identified reduced transcripts and ribosome occupancy in pathways involving ion homeostasis, potassium channels, and cardiac conduction, while translation-, RNA processing-, and ribosome-related pathways emerged (Figure 4A–D). This suggests a transition from an early inflammatory/stress state toward a later phase marked by persistent translational remodeling and disruption of pathways necessary for normal cardiac function.

**Figure 4.**
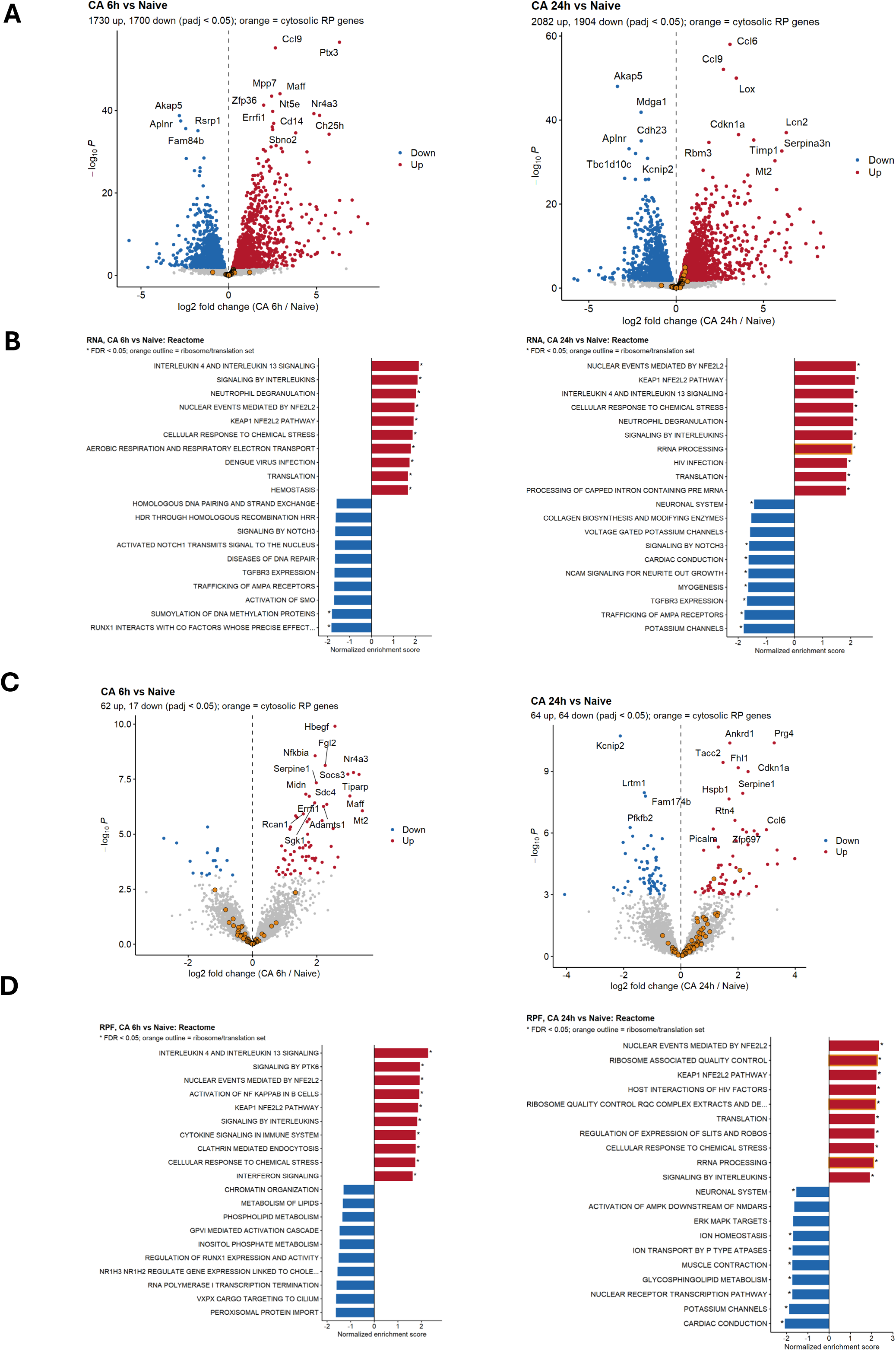
Transcript abundance and ribosome occupancy after cardiac arrest. A) Volcano plots of RNA-seq comparing CA 6h (left) and CA 24h (right) with Naïve controls; cytosolic ribosomal protein (RP) genes are highlighted in orange. B) Top 10 up- and downregulated Reactome pathways by gene set enrichment analysis of RNA-seq at 6 h (left) and 24 h (right); ribosome and translation pathways are outlined in orange. C) Volcano plots of Ribo-seq for the same comparisons. D) Top 10 up- and downregulated Reactome pathways by gene set enrichment analysis of Ribo-seq at 6 h (left) and 24 h (right). n = 6 Naïve, 6 CA 6h, 5 CA 24h. Differential expression by DESeq2 with sex and sequencing batch as covariates; adjusted p < 0.05 by Benjamini-Hochberg. *=FDR < 0.05 for pathway enrichment

At 6 h, no pathways reached significance by TE analysis (Figure 5A). By 24 h, ribosome- and translation-related pathways, including eukaryotic translation initiation and elongation, ribosome-associated quality control, and nonsense-mediated decay, showed the strongest positive enrichment by TE (Figure 5A). This suggests that ribosome occupancy shifted toward mRNAs encoding the translational machinery despite reduced global protein synthesis. Proteomic analysis showed similar changes to translation associated products. Cytosolic RP proteins were shifted toward increased abundance at both time points (Figure 5B), and translation, ribosome quality control, and translation initiation and elongation pathways were enriched at 6 h and 24 h (Figure 5C). Mitochondrial translation pathways were also enriched at 6 h. Across all modalities and time points, translation, ribosome quality control, and RNA processing pathways were among the most consistently enriched programs, most prominently at 24 h (Figure 6).

**Figure 5.**
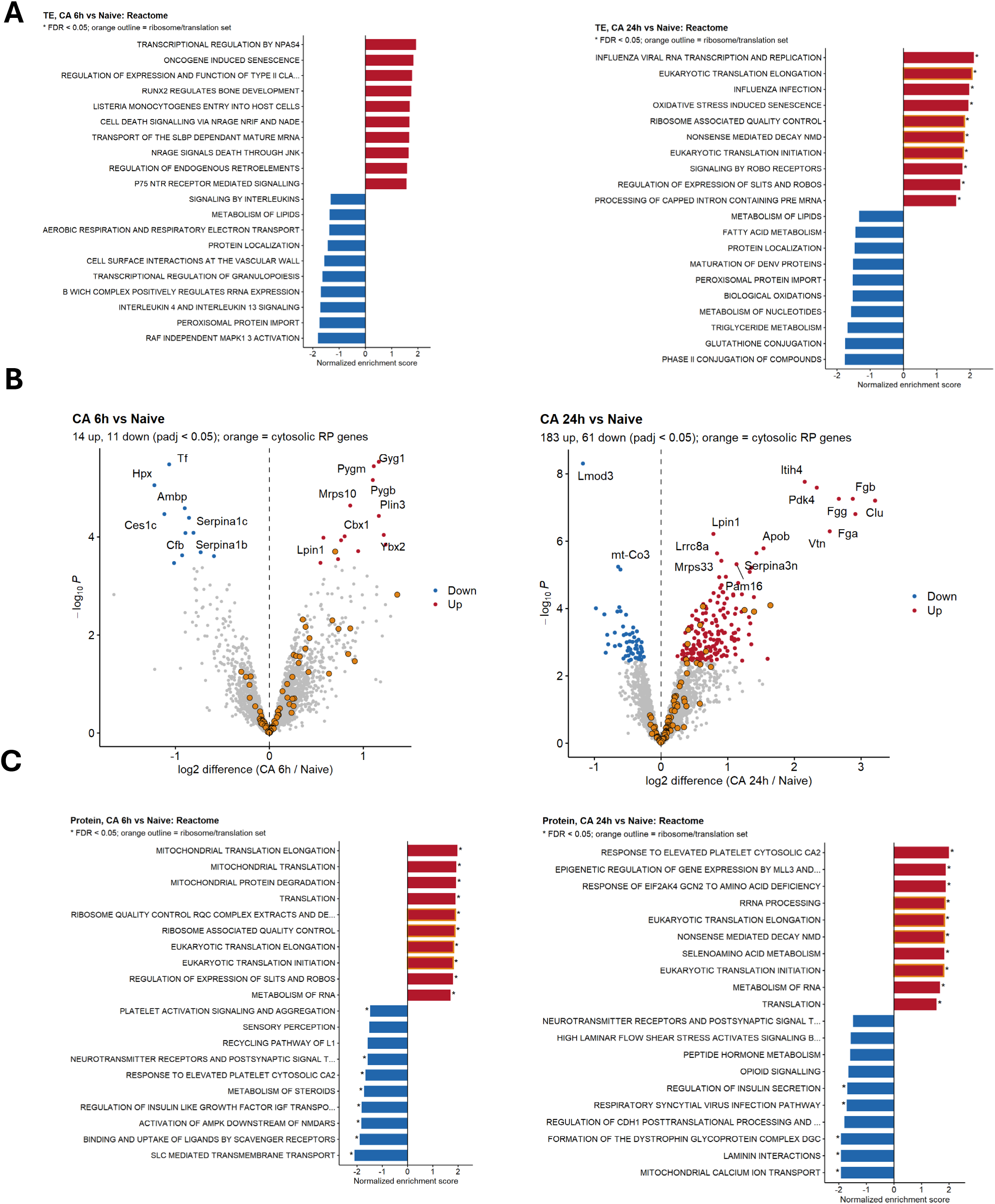
Translational efficiency and protein-level pathway responses after cardiac arrest. A) Top 10 up- and downregulated pathways by gene set enrichment analysis of translational efficiency at 6 h (left) and 24 h (right); ribosome and translation pathways are outlined in orange. B) Volcano plots of proteomics comparing CA 6h (left) and CA 24h (right) with Naïve controls; cytosolic RP proteins are highlighted in orange. C) Top 10 up- and downregulated Reactome pathways by gene set enrichment analysis of proteomics at 6 h (left) and 24 h (right). n = 6 Naïve, 6 CA 6h, 5 CA 24h for translational efficiency; n = 6 Naïve, 4 CA 6h, 6 CA 24h for proteomics. Translational efficiency by deltaTE; proteomics by limma with sex as a covariate; adjusted p < 0.05 by Benjamini-Hochberg. *=FDR < 0.05 for pathway enrichment.

**Figure 6.**
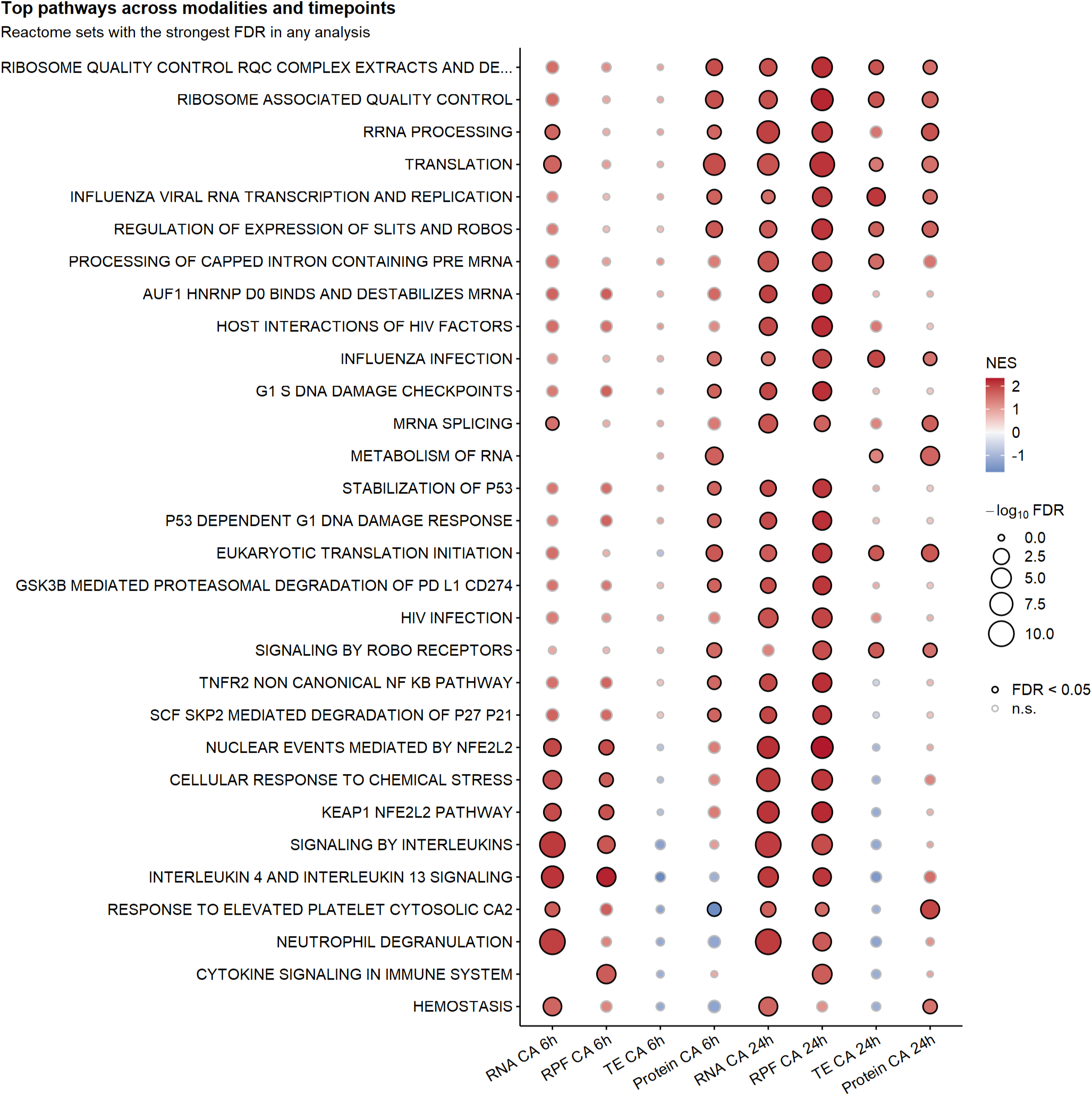
Summary of pathway analysis across modalities and timepoints. Dot plot of the Reactome gene sets with the strongest FDR in any analysis, compared across RNA-seq (RNA), ribosome profiling (RPF), translational efficiency (TE), and proteomics (Protein) at 6 h and 24 h after cardiac arrest (CA) versus Naïve controls. Rows are pathways. Columns are modality and time point. Dot color indicates the normalized enrichment score (NES) from gene set enrichment analysis (red, positive; blue, negative). Dot size indicates −log₁₀ FDR. Dots with a black outline have FDR < 0.05, and faint dots without an outline are not significant (n.s.). Empty positions indicate that the pathway was not tested in that analysis. n = 6 Naïve, 6 CA 6h, and 5 CA 24h for RNA, RPF, and TE; n = 6 Naïve, 4 CA 6h, and 6 CA 24h for proteomics. Differential expression was analyzed by DESeq2 (RNA, RPF), deltaTE (TE), and limma (Protein); adjusted p < 0.05 by Benjamini-Hochberg.

Inflammatory pathways induced at the RNA level at 6 h were attenuated or absent at the TE and protein levels. Collectively, these findings indicate that CA remodels myocardial gene expression at multiple regulatory levels. As global protein synthesis declines, ribosome occupancy and protein abundance shift toward the translational machinery to compensate.

## Discussion

Myocardial dysfunction following CA has no targeted therapies and current treatment strategies remain largely supportive. While many investigative therapies in CA are targeting specific inflammatory or signaling molecules, broad and more integrated mechanisms of injury should be considered as potential targets. In this study, we identify ribosomal remodeling and translational suppression as central features of the myocardial response to CA. Across complementary porcine and murine models, we demonstrate that CA increases ribosomal transcript abundance, and that increasing ischemic burden is associated with more pronounced response in the pig model. However, direct assessment of protein synthesis in the mouse model demonstrates global suppression of myocardial translation rates, and that this suppression lasts for at least 24 hours after resuscitation. Multi-omic profiling in the mouse model demonstrates that transcript abundance, ribosome occupancy, TE, and protein level response change dynamically after CA, but that ribosome-associated pathways change most dramatically, outpacing inflammatory targets. Increased ribosomal transcripts should not be interpreted as evidence of increased ribosome activity, but rather as a compensatory response to restore or rebuild translational machinery after IRI.

This compensatory response is consistent with prior literature demonstrating translational suppression in response to IRI across multiple tissues^18,22,27,37^. Decreased ribosome activity in response to hypoxia is generally thought to represent an adaptive response to conserve energy and promote cell survival^17,19,38^. However, persistent translational suppression after reperfusion may be maladaptive if it impairs replacement of damaged proteins or impairs the ability to restore critical organ functions^38^. Similar studies of IRI in the brain have found that reduced ribosome activity can lead to neuronal death in specific regions after IRI^22,39^. In the heart, inability to restore ribosomal machinery may contribute to inability to maintain critical functions including ion homeostasis, mitochondrial integrity, conduction and contraction.

Multi-omic analyses of the mouse heart support the framework of prolonged suppression after reperfusion, and also help to characterize the consequences of this translational deficit. TE plots demonstrate high correlation between ribosome-associated transcripts in Naïve and both 6 h and 24 h cohorts (Figure 3C), suggesting that translation is suppressed globally across transcripts. However, profiling of these ribosome-associated changes in ribosome demonstrates upregulation of inflammatory and stress-response pathways in the 6 h cohort compared to Naïve, with no significantly downregulated responses. By 24 h, downregulation of pathways related to ion homeostasis, potassium channels, and cardiac conduction become more evident by ribosome profiling, while upregulated pathways reflect translation of ribosome-related pathways, likely as a compensatory response, and to a lesser extent, inflammatory pathways. Impaired translation may contribute directly to cardiac dysfunction by limiting nascent proteins contributing to contractility and ion handling.

Proteome data offer limited insight to translational changes. Proteins have long half-lives and proteome detection is limited to the most abundant proteins, so it is difficult to obtain the depth of insights available by RNA or ribosome sequencing. Further, proteins are actively degraded after the insult, introducing complications related to autophagy and protein clearance. Despite these limitations, proteome data does highlight notable changes in the post-arrest heart, including increased signaling related to both cytoplasmic and mitochondrial translation and ribosome control, altered calcium handling pathways, and metabolic signaling related to insulin secretion and AMPK signaling (Figure 5C).

The implication of ribosome signaling in both models and across multiple time-points and -omic modalities raise the possibility of therapeutic options aimed at restoring ribosome function. Prior work in our lab has shown that metformin pretreatment preserves ribosome activity after CA, but does not work as a rescue therapy, and metformin’s protective effects are supported by clinical data^27,40^. Future studies are warranted to explore pharmacologic and nutritional strategies that may preserve ribosome activity, which may offer a broader and potentially more effective strategy than targeting individual downstream pathways.

Several limitations should be acknowledged. First, RNA integrity is a concern in ischemic tissue, and degradation biases transcript quantification in a gene-specific manner that may favor short, abundant transcripts such as those encoding ribosomal protein^41,42^. However, RIN was included as a covariate in all porcine analyses, RP transcript abundance did not correlate with RIN (Supplemental Figure 2C), and ribosomal pathways were also enriched in murine myocardium, where RNA integrity was uniformly high. Second, the porcine and murine models cannot fully capture the heterogeneity of human CA. In pigs, remote myocardium rather than border-zone tissue was sampled in order to minimize confounding effects of tissue necrosis, though remote tissue may underestimate local injury adjacent to the infarct zone. In mice, KCl-induced asystole is necessary in order to study the mouse model, which enables longer term and puromycin incorporation studies, but may not reflect the mechanisms or severity of clinical CA. Third, the puromycin incorporation studies provide a functional assay of nascent protein synthesis, but uptake my be influenced by local reperfusion or altered drug metabolism in the CA model and may not fully recapitulate ribosome activity. Finally, this study focused on cytoplasmic ribosome changes and global translation, but does not measure mitochondrial-specific translation, which may be an alternative mechanism of post-arrest recovery.

In conclusion, CA induces marked remodeling of myocardial ribosome biology. Ribosomal transcripts increase with ischemic burden, while protein synthesis is suppressed for at least 24 h after resuscitation, coinciding with reduced ribosome occupancy on transcripts required for conduction, ion homeostasis, and contraction. These findings identify translational control as a central feature of post-arrest myocardial injury and suggest that restoring ribosome function may be a novel therapeutic strategy for post-arrest cardiac dysfunction.

## Abbreviations

ACLS: Advanced Cardiac Life Support
ANOVA: Analysis of Variance
CA: Cardiac Arrest
CPR: Cardiopulmonary resuscitation
DEG: Differentially Expressed Gene
EF: Ejection Fraction
FDR: False Discovery Rate
GSEA: Gene Set Enrichment Analysis
IRI: Ischemia Reperfusion Injury
KCl: Potassium Chloride
LAD: Left Anterior Descending
LV: Left Ventricle
MAP: Mean Arterial Pressure
MI: Myocardial Infarction
NES: Normalized Enrichment Score
PCA: Principal Component Analysis
PEA: Pulseless Electrical Activity
Ribo-seq: Ribosome Sequencing
RIN: RNA Integrity Number
RP: Ribosomal Protein
RPF: Ribosome-Protected Fragment
ROSC: Return of Spontaneous Circulation
SEM: Standard Error of the Mean
TBS: Tris Buffered Saline
TE: Translational Efficiency
VF: Ventricular Fibrillation

## Contributions

KL, LW, and CR were responsible for the conceptualization of these studies. YG, WD, KL, LW and CR designed the methodology and performed the investigation. Formal analyses were performed by GZ, KL, LW, ZG, RX and CR. GZ, ZG, RX, and CR completed visualization. YG, KL, LW, and CR wrote the manuscript. CR supervised the project and provided resources for its completion. All authors reviewed the final manuscript and are responsible for its integrity.

## References

1. Palaniappan LP, Allen NB, Almarzooq ZI, et al. 2026 Heart Disease and Stroke Statistics: A Report of US and Global Data From the American Heart Association. Circulation. 2026;153(9):e275–e906. doi:10.1161/CIR.0000000000001412

2. 2025 CARES Annual Report. Accessed October 13, 2026. https://mycares.net/sitepages/uploads/2026/2025_flipbook/index.html?page=28

3. Committee on the Treatment of Cardiac Arrest: Current Status and Future Directions, Board on Health Sciences Policy, Institute of Medicine. Strategies to Improve Cardiac Arrest Survival: A Time to Act. (Graham R, McCoy MA, Schultz AM, eds.). National Academies Press (US); 2015. Accessed November 5, 2026. http://www.ncbi.nlm.nih.gov/books/NBK305685/

4. Laver S, Farrow C, Turner D, Nolan J. Mode of death after admission to an intensive care unit following cardiac arrest. Intensive Care Med. 2004;30(11):2126–2128. doi:10.1007/s00134-004-2425-z

5. Jozwiak M, Bougouin W, Geri G, Grimaldi D, Cariou A. Post-resuscitation shock: recent advances in pathophysiology and treatment. Ann Intensive Care. 2020;10:170. doi:10.1186/s13613-020-00788-z

6. Neumar RW, Nolan JP, Adrie C, et al. Post–Cardiac Arrest Syndrome: Epidemiology, Pathophysiology, Treatment, and Prognostication A Consensus Statement From the International Liaison Committee on Resuscitation (American Heart Association, Australian and New Zealand Council on Resuscitation, European Resuscitation Council, Heart and Stroke Foundation of Canada, InterAmerican Heart Foundation, Resuscitation Council of Asia, and the Resuscitation Council of Southern Africa); the American Heart Association Emergency Cardiovascular Care Committee; the Council on Cardiovascular Surgery and Anesthesia; the Council on Cardiopulmonary, Perioperative, and Critical Care; the Council on Clinical Cardiology; and the Stroke Council. Circulation. 2008;118(23):2452–2483. doi:10.1161/CIRCULATIONAHA.108.190652

7. Lemiale V, Dumas F, Mongardon N, et al. Intensive care unit mortality after cardiac arrest: the relative contribution of shock and brain injury in a large cohort. Intensive Care Med. 2013;39(11):1972–1980. doi:10.1007/s00134-013-3043-4

8. Ruiz-Bailén M, Hoyos EAD, Ruiz-Navarro S, et al. Reversible myocardial dysfunction after cardiopulmonary resuscitation. Resuscitation. 2005;66(2):175–181. doi:10.1016/j.resuscitation.2005.01.012

9. Laurent I, Monchi M, Chiche JD, et al. Reversible myocardial dysfunction in survivors of out-of-hospital cardiac arrest. Journal of the American College of Cardiology. 2002;40(12):2110–2116. doi:10.1016/S0735-1097(02)02594-9

10. Kern KB, Hilwig RW, Rhee KH, Berg RA. Myocardial dysfunction after resuscitation from cardiac arrest: An example of global myocardial stunning. Journal of the American College of Cardiology. 1996;28(1):232–240. doi:10.1016/0735-1097(96)00130-1

11. Heusch G. Myocardial stunning and hibernation revisited. Nat Rev Cardiol. 2021;18(7):522–536. doi:10.1038/s41569-021-00506-7

12. Ibáñez B, Heusch G, Ovize M, Van de Werf F. Evolving Therapies for Myocardial Ischemia/Reperfusion Injury. Journal of the American College of Cardiology. 2015;65(14):1454–1471. doi:10.1016/j.jacc.2015.02.032

13. Hirsch KG, Amorim E, Coppler PJ, et al. Part 11: Post-Cardiac Arrest Care: 2025 American Heart Association Guidelines for Cardiopulmonary Resuscitation and Emergency Cardiovascular Care. Circulation. 2025;152(16_suppl_2):S673–S718. doi:10.1161/CIR.0000000000001375

14. Meyer MAS, Wiberg S, Grand J, et al. Treatment Effects of Interleukin-6 Receptor Antibodies for Modulating the Systemic Inflammatory Response After Out-of-Hospital Cardiac Arrest (The IMICA Trial): A Double-Blinded, Placebo-Controlled, Single-Center, Randomized, Clinical Trial. Circulation. 2021;143(19):1841–1851. doi:10.1161/CIRCULATIONAHA.120.053318

15. Argaud L, Cour M, Dubien PY, et al. Effect of Cyclosporine in Nonshockable Out-of-Hospital Cardiac Arrest: The CYRUS Randomized Clinical Trial. JAMA Cardiol. 2016;1(5):557–565. doi:10.1001/jamacardio.2016.1701

16. Davidson SM, Ferdinandy P, Andreadou I, et al. Multitarget Strategies to Reduce Myocardial Ischemia/Reperfusion Injury: JACC Review Topic of the Week. J Am Coll Cardiol. 2019;73(1):89–99. doi:10.1016/j.jacc.2018.09.086

17. Liu L, Cash TP, Jones RG, Keith B, Thompson CB, Simon MC. Hypoxia-Induced Energy Stress Regulates mRNA Translation and Cell Growth. Molecular Cell. 2006;21(4):521–531. doi:10.1016/j.molcel.2006.01.010

18. DeGracia DJ, Kumar R, Owen CR, Krause GS, White BC. Molecular Pathways of Protein Synthesis Inhibition during Brain Reperfusion: Implications for Neuronal Survival or Death. J Cereb Blood Flow Metab. 2002;22(2):127–141. doi:10.1097/00004647-200202000-00001

19. Fähling M. Surviving hypoxia by modulation of mRNA translation rate. J Cellular Molecular Medi. 2009;13(9a):2770–2779. doi:10.1111/j.1582-4934.2009.00875.x

20. Hossmann KA. Disturbances of cerebral protein synthesis and ischemic cell death. Prog Brain Res. 1993;96:161–177. doi:10.1016/s0079-6123(08)63265-3

21. Simon MC, Liu L, Barnhart BC, Young RM. Hypoxia-Induced Signaling in the Cardiovascular System. Annu Rev Physiol. 2008;70:51–71. doi:10.1146/annurev.physiol.70.113006.100526

22. Krause GS, Tiffany BR. Suppression of protein synthesis in the reperfused brain. Stroke. 1993;24(5):747–755. doi:10.1161/01.STR.24.5.747

23. DeGracia DJ. Acute and persistent protein synthesis inhibition following cerebral reperfusion. J Neurosci Res. 2004;77(6):771–776. doi:10.1002/jnr.20225

24. Lewis YE, Moskovitz A, Mutlak M, Heineke J, Caspi LH, Kehat I. Localization of transcripts, translation, and degradation for spatiotemporal sarcomere maintenance. J Mol Cell Cardiol. 2018;116:16–28. doi:10.1016/j.yjmcc.2018.01.012

25. Kim TY, Wang D, Kim AK, et al. Metabolic labeling reveals proteome dynamics of mouse mitochondria. Mol Cell Proteomics. 2012;11(12):1586–1594. doi:10.1074/mcp.M112.021162

26. Balse E, Steele DF, Abriel H, Coulombe A, Fedida D, Hatem SN. Dynamic of ion channel expression at the plasma membrane of cardiomyocytes. Physiol Rev. 2012;92(3):1317–1358. doi:10.1152/physrev.00041.2011

27. Rutledge CA, Lagranha C, Chiba T, et al. Metformin preconditioning protects against myocardial stunning and preserves protein translation in a mouse model of cardiac arrest. Journal of Molecular and Cellular Cardiology Plus. 2023;4:100034. doi:10.1016/j.jmccpl.2023.100034

28. Piktel JS, Suen Y, Kouk S, et al. Effect of Amiodarone and Hypothermia on Arrhythmia Substrates During Resuscitation. J Am Heart Assoc. 2021;10(10):e016676. doi:10.1161/JAHA.120.016676

29. Laurita KR, Piktel JS, Irish L, et al. Spontaneous Repolarization Alternans Causes VT/VF Rearrest That Is Suppressed by Preserving Gap Junctions. JACC Clin Electrophysiol. 2024;10(7 Pt 1):1271–1286. doi:10.1016/j.jacep.2024.03.027

30. Rutledge CA, Chiba T, Redding K, Dezfulian C, Sims-Lucas S, Kaufman BA. A novel ultrasound-guided mouse model of sudden cardiac arrest. Johnson DM, ed. PLoS ONE. 2020;15(12):e0237292. doi:10.1371/journal.pone.0237292

31. Rutledge C, Cater G, McMahon B, et al. Commercial 4-dimensional echocardiography for murine heart volumetric evaluation after myocardial infarction. Cardiovasc Ultrasound. 2020;18(1):9. doi:10.1186/s12947-020-00191-5

32. Abella BS, Zhao D, Alvarado J, Hamann K, Vanden Hoek TL, Becker LB. Intra-Arrest Cooling Improves Outcomes in a Murine Cardiac Arrest Model. Circulation. 2004;109(22):2786–2791. doi:10.1161/01.CIR.0000131940.19833.85

33. Schmidt EK, Clavarino G, Ceppi M, Pierre P. SUnSET, a nonradioactive method to monitor protein synthesis. Nat Methods. 2009;6(4):275–277. doi:10.1038/nmeth.1314

34. Schindelin J, Arganda-Carreras I, Frise E, et al. Fiji: an open-source platform for biological-image analysis. Nat Methods. 2012;9(7):676–682. doi:10.1038/nmeth.2019

35. Rao A, Lyu B, Jahan I, et al. The translation initiation factor homolog eif4e1c regulates cardiomyocyte metabolism and proliferation during heart regeneration. Development. 2023;150(20):dev201376. doi:10.1242/dev.201376

36. Ingolia NT, Ghaemmaghami S, Newman JRS, Weissman JS. Genome-wide analysis in vivo of translation with nucleotide resolution using ribosome profiling. Science. 2009;324(5924):218–223. doi:10.1126/science.1168978

37. Montie HL, Kayali F, Haezebrouck AJ, Rossi NF, DeGracia DJ. Renal ischemia and reperfusion activates the eIF2 alpha kinase PERK. Biochimica et Biophysica Acta (BBA) - Molecular Basis of Disease. 2005;1741(3):314–324. doi:10.1016/j.bbadis.2005.04.007

38. Kidwell A, Yadav SPS, Maier B, et al. Translation Rescue by Targeting Ppp1r15a through Its Upstream Open Reading Frame in Sepsis-Induced Acute Kidney Injury in a Murine Model. Journal of the American Society of Nephrology.: JASN. 2022;34(2):220. doi:10.1681/ASN.2022060644

39. DeGracia DJ, Hu BR. Irreversible translation arrest in the reperfused brain. Journal of cerebral blood flow and metabolism.: official journal of the International Society of Cerebral Blood Flow and Metabolism. 2006;27(5):875. doi:10.1038/sj.jcbfm.9600388

40. Rutledge CA, Kaufman BA, Dezfulian C, Elmer J. Metformin protects against cardiac and renal damage in diabetic cardiac arrest patients. Resuscitation. 2022;174:42–46. doi:10.1016/j.resuscitation.2022.03.017

41. Gallego Romero I, Pai AA, Tung J, Gilad Y. RNA-seq: impact of RNA degradation on transcript quantification. BMC Biol. 2014;12:42. doi:10.1186/1741-7007-12-42

42. Xiong B, Yang Y, Fineis FR, Wang JP. DegNorm: normalization of generalized transcript degradation improves accuracy in RNA-seq analysis. Genome Biol. 2019;20:75. doi:10.1186/s13059-019-1682-7

