## Supplemental Figures for "Ribosomal Remodeling and Translational Suppression in the Post-Arrest Heart"

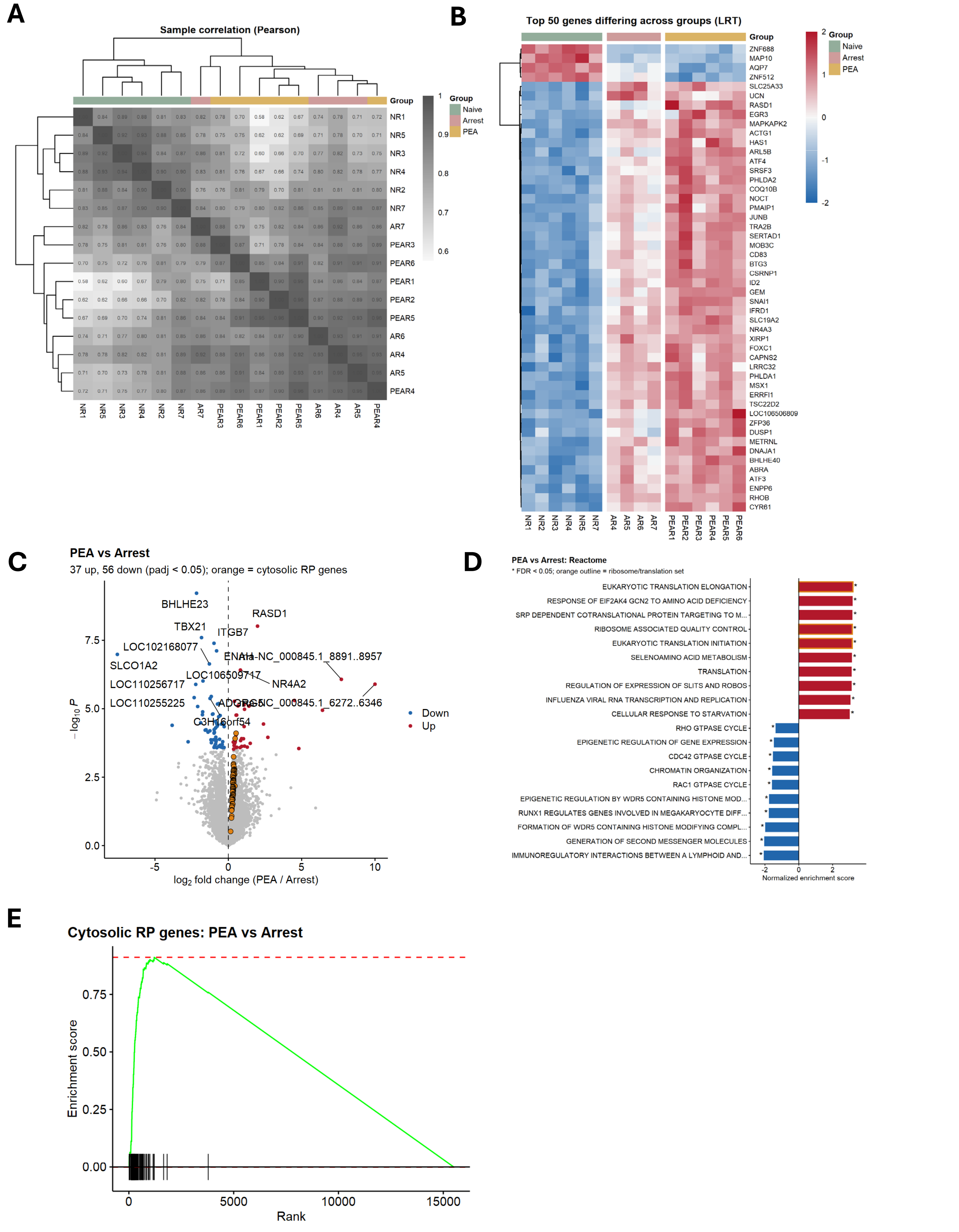


**Supplemental Figure 1. Transcriptomic separation by group and direct comparison of PEA to Arrest animals.** A) Pearson correlation matrix of remote myocardial transcriptomes, clustered by 1 – correlation distance. B) Heatmap of the 50 genes most significantly differing across groups by likelihood ratio test, shown as row z-scores. C) Volcano plot comparing PEA to Arrest animals, with cytosolic RP genes highlighted in orange. D) Top 10 up- and downregulated Reactome pathways for PEA versus Arrest; ribosome and translation pathways are outlined in orange. E) Enrichment plot for the 78 cytosolic RP genes in PEA versus Arrest. n = 6 Naïve, 4 Arrest, 6 PEA. *=FDR < 0.05

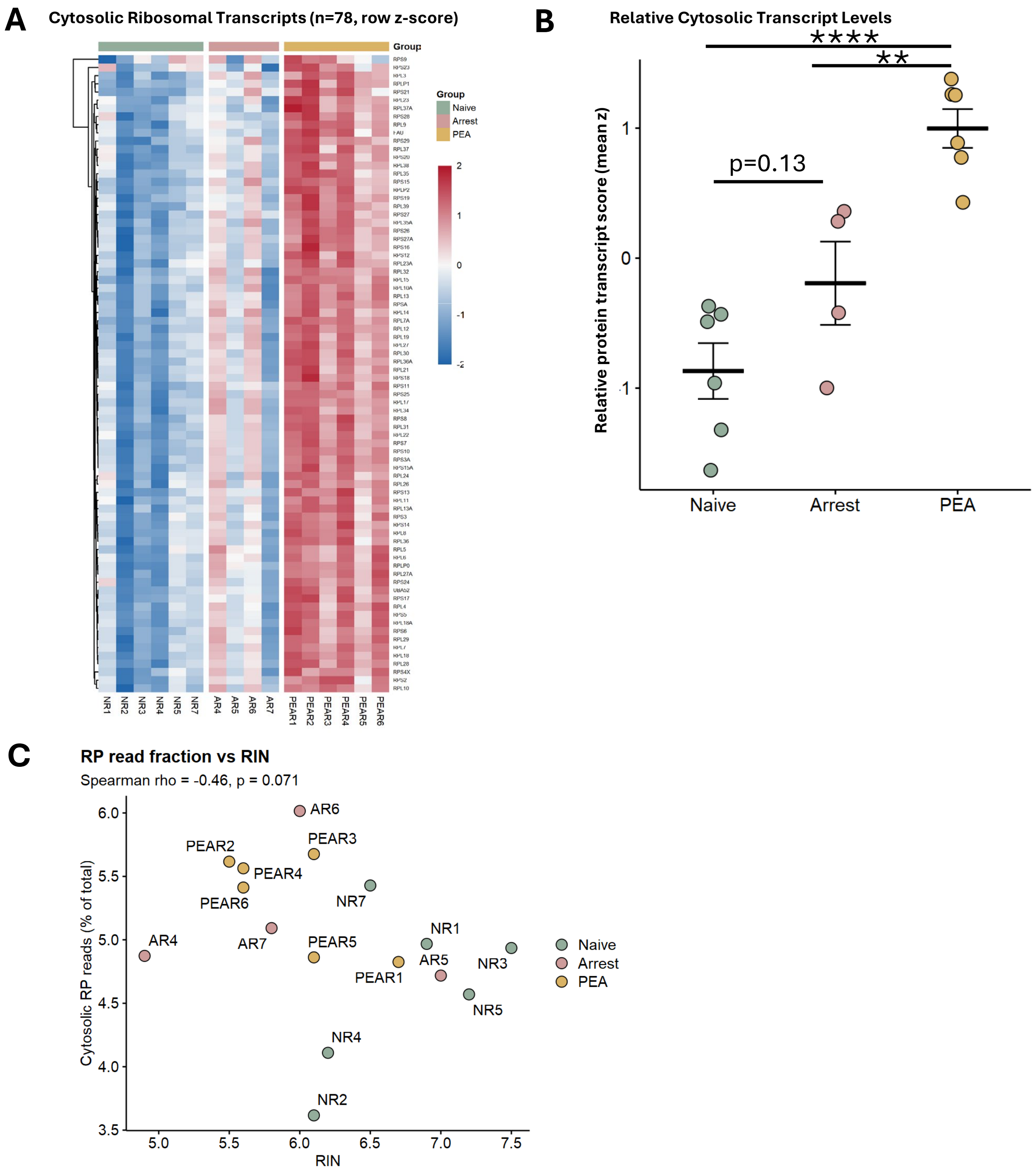


**Supplemental Figure 2. Ribosomal protein expression increases with ischemic burden and not explained by RNA integrity.** A) Heatmap of the 78 detected cytosolic RP genes, shown as row z-scores with samples ordered by group. B) Per-sample RP score, calculated as the mean z-score of these genes after adjustment for RIN and sequencing run. C) Fraction of total reads assigned to cytosolic RP genes plotted against RIN for each sample. n = 6 Naïve, 4 Arrest, 6 PEA. Data expressed as mean ± SEM. **=p<0.01, ****=p<0.0001 by one way ANOVA.

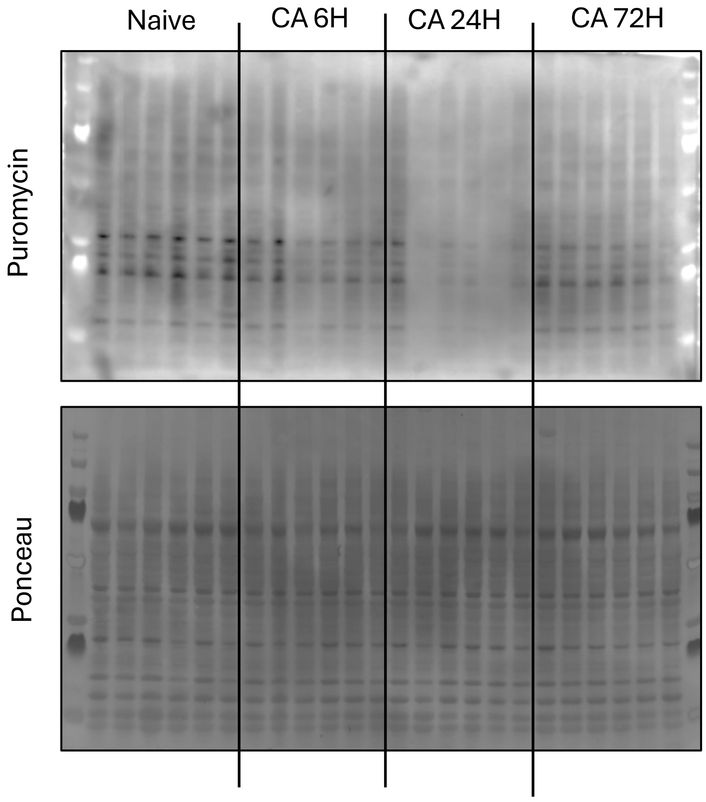


**Supplemental Figure 3. Full puromycin immunoblot.** Full-length puromycin immunoblot (top) and corresponding Ponceau S total protein stain (bottom) of left ventricular lysates from Naïve, CA 6 h, CA 24 h, and CA 72 h mice. Each lane is an individual animal, and molecular weight markers flank the samples. All samples were run on a single gel and transferred to a single membrane. These blots were quantified in Figure 2C. n = 6/group.

A

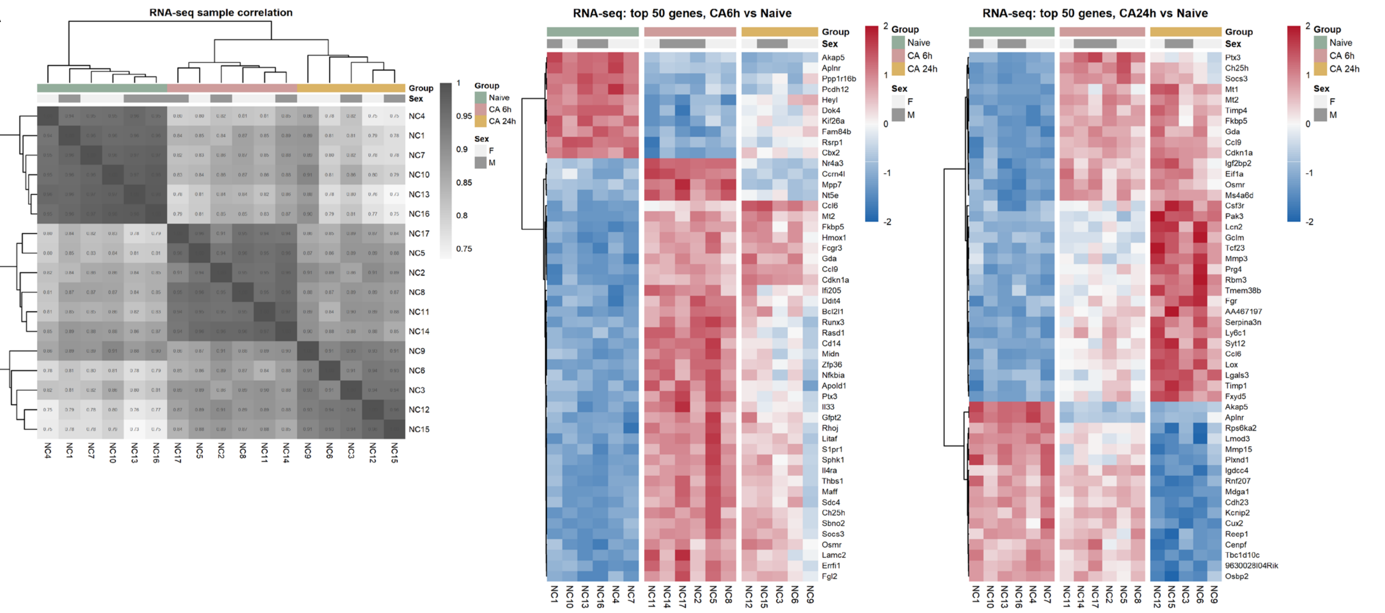


**Supplemental Figure 4. RNA-seq sample clustering and differential expression.** A) Pearson correlation matrix of RNA-seq samples, clustered by 1 − correlation distance (left), and heatmaps of the 50 most significantly changed genes at 6 h (center) and 24 h (right), shown as row z-scores. n = 6 Naïve, 6 CA 6h, 5 CA 24h.

A

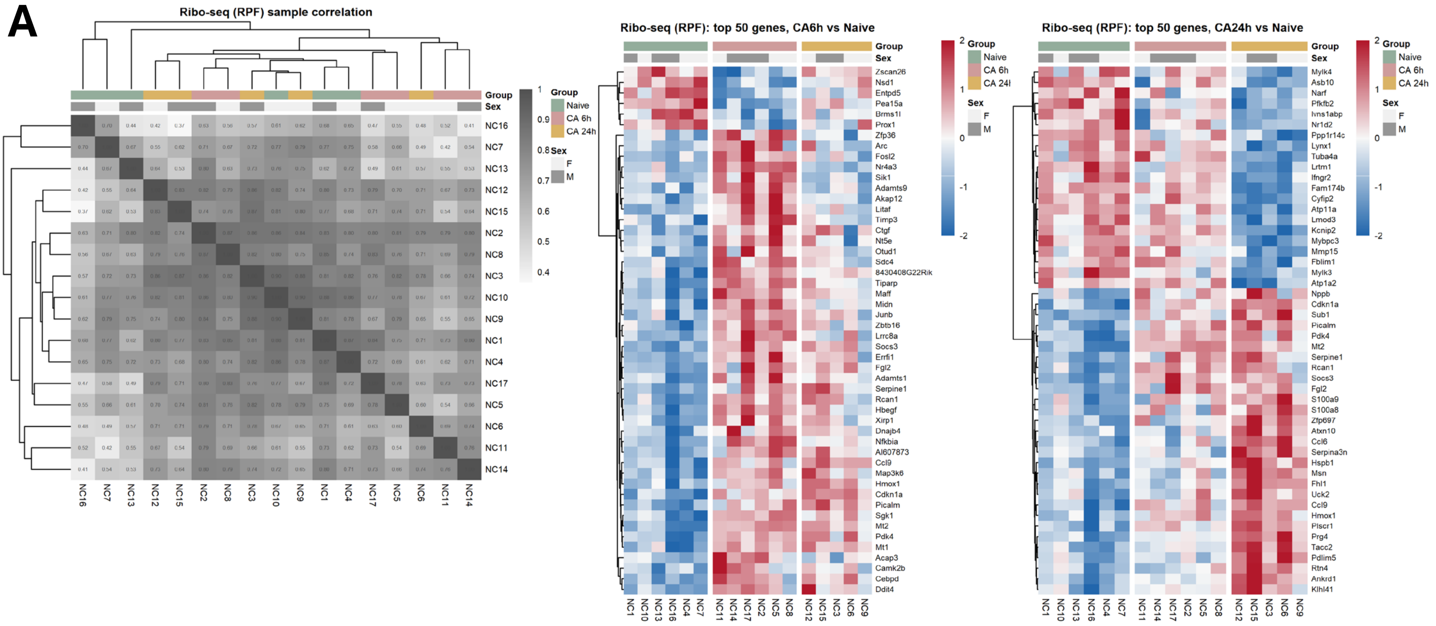

**Supplemental Figure 5. Ribo-seq sample clustering and differential expression.** A) Pearson correlation matrix of ribosome profiling samples, clustered by 1 − correlation distance (left), and heatmaps of the 50 most significantly changed genes at 6 h (center) and 24 h (right), shown as row z-scores. n = 6 Naïve, 6 CA 6h, 5 CA 24h

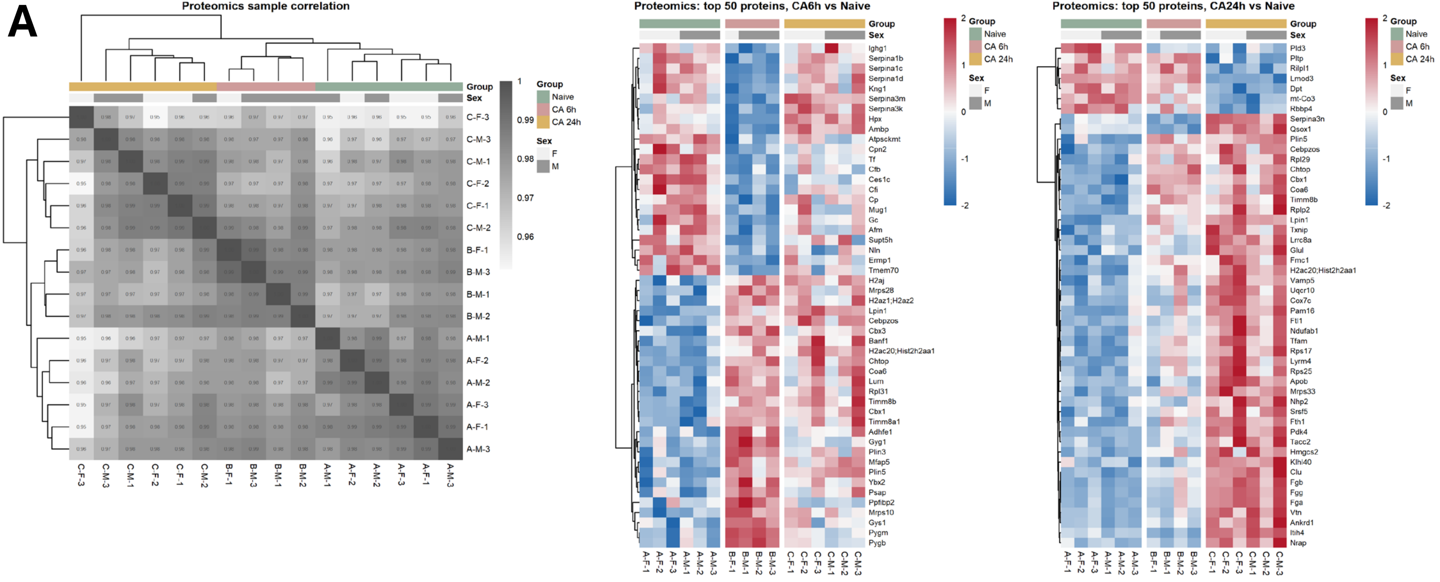


**Supplemental Figure 6. Proteomic sample clustering and differential abundance.** A) Pearson correlation matrix of proteomic samples, clustered by 1 − correlation distance (left), and heatmaps of the 50 most significantly changed proteins at 6 h (center) and 24 h (right), shown as row z-scores. n = 6 Naïve, 4 CA 6h, 6 CA 24h.
